# Cryo-EM structure of a cell-free synthesized full-length human β1-adrenergic receptor in complex with G_s_

**DOI:** 10.1101/2025.04.29.651181

**Authors:** Felipe Merino, Zoe Köck, Utz Ermel, Philipp Dahlhaus, Anna Grimm, Anja Seybert, Jan Kubicek, Achilleas S. Frangakis, Volker Dötsch, Daniel Hilger, Frank Bernhard

## Abstract

The third intracellular loop (ICL3) of the β1-adrenergic receptor (β1AR) plays a critical role in regulating G protein coupling, yet the structural basis has remained unclear due to truncations of ICL3 in all available structures of the β1AR in complex with G_s_ or a G protein mimetic nanobody. To address this, we used cell-free cotranslational insertion of full-length human β1AR into nanodiscs and determined its cryo-EM structure in complex with G_s_. In this structure, ICL3 extends transmembrane helix 5, resulting in enhanced interactions with Gα_s_ and in a slight rotation of the engaged G protein. This repositioning enables new polar interactions between Gα_s_, ICL2 and helix 8, while ICL1 and helix 8 form additional contacts with Gβ. These structural insights, supported by mutational analysis, demonstrate that ICL3 enhances G protein activation and downstream cAMP signaling by promoting more extensive interactions between the receptor and the heterotrimeric G protein.

## INTRODUCTION

G protein coupled receptors (GPCRs) play pivotal roles in treating many human diseases, therefore being key targets for pharmaceutical research.^1^ Recombinant GPCR expression is an essential prerequisite for functional and structural analysis. In conventional eukaryotic cell-based expression systems, GPCRs must pass various membrane systems during their transport to the cell surface. Critical steps such as translocon-assisted membrane insertion, limited membrane space, and incorrect glycosylation or trafficking mechanisms often limit recombinant GPCR synthesis.^2, 3^ To overcome limitations by proteolysis or low surface expression, engineering strategies such as deleting the unstructured terminal domains and truncating the flexible intracellular loop 3 (ICL3) are commonly applied.^4–9^ Additionally, chimeric proteins are often generated by inserting well-folding domains, such as T4 lysozyme or thermostabilized apocytochrome b562 (BRIL), to stabilize recombinant GPCRs for downstream processing and structural characterization.^10^ While these modifications generally preserve the basic functionality of engineered GPCRs, they can result in local structural distortions and affect conformational flexibility.^11^ Furthermore, the drastic alteration of the hydrophobic environment during detergent extraction of recombinant GPCRs from cell membranes may lead to significant protein destabilization and denaturation.^12^

The cotranslational insertion of GPCRs into defined lipid environments of nanodiscs (NDs) by cell-free (CF) expression avoids any exposure to detergents and artificial reconstitution procedures.^13–16^ This reduces the risk of GPCR denaturation, facilitates their characterization in different lipid environments, and enables the study of detergent-sensitive targets.^17^ Furthermore, the controlled addition of ligands or interacting proteins at stoichiometric concentrations during CF translation can already stabilize the nascent GPCRs, enabling the production of full-length GPCRs with highly flexible domains, such as the human β1-adrenergic receptor (Hβ1AR).^16, 18^ These benefits recently resulted into the cryogenic-electron microscopy (cryo-EM) structure of the CF synthesized full-length human histamine 2 receptor in complex with G_s_ heterotrimer.^19^

The β1AR is one of the best-characterized class A GPCRs, the largest and most diverse GPCR subfamily in humans. It plays a crucial role in regulating heart rate and contractility in response to catecholamines such as noradrenaline and adrenaline.^20^ Numerous atomic structures of β1AR have been determined in complex with various agonists or antagonists, and coupled to its cognate heterotrimeric G proteins, G_s_ and G_i_, and β-arrestin.^21–26^ Due to its higher stability in detergent, all available β1AR-G protein complex structures have been solved using a truncated turkey ortholog (TΔβ1AR).^27^ For Hβ1AR, no structures in complex with G proteins have been determined so far and only four active state structures are available, stabilized by the G protein mimetic intracellular nanobody.^10^ Similar to its turkey ortholog, successful structural characterization of Hβ1AR required truncations of the N- and C-termini and ICL3 to produce well-diffracting crystals (hereby referred to as HΔβ1AR). Notably, ICL3 has been shown to play a role in efficient G protein coupling and cAMP signaling, likely by enhancing the interaction between the receptor and the Gαs subunit following GDP release.^28^ However, the structural basis of the ICL3 impact on G_s_ coupling to the full-length Hβ1AR remains poorly understood.

Here, we combine CF expression and cryo-EM to determine the structure of a thermostabilized full-length Hβ1AR (HFLβ1AR) bound to isoprenaline and in complex with its canonical heterotrimeric G protein G_s_ in NDs. Combined with mutagenesis and signaling studies, our structural analysis reveals that the presence of ICL3 significantly improves G protein coupling through formation of additional contacts between HFLβ1AR and the Gα_s_ and Gβ subunit of the heterotrimeric G_s_ protein ultimately leading to enhanced cAMP signaling.

## RESULTS

### Cryo-EM structure of the isoprenaline-bound HFLβ1AR-G_s_ complex in nanodiscs

To prepare a stable HFLβ1AR-G_s_ complex, we synthesized the HFLβ1AR G389^8×56^ variant (amino acids 1-477) via CF expression in *E. coli* A19 S30 lysates (the superscript numbers refer to the generic GPCR numbering based on the revised Ballesteros-Weinstein system for class A GPCRs).^29, 30^ The receptor was cotranslationally inserted into preformed NDs containing the lipid 1,2-dioleoyl-sn*-*glycero-3-phosphoglycerol (DOPG), enabling the efficient membrane insertion and folding of HFLβ1AR.^16^ Nine point mutations – K85^1×59^S, M107^2×53^V, I146^3×40^V, E147^3×41^W, A316^6×27^L, F361^7×36^A, F372^7×48^M, C392^8×59^A and C393A – were introduced to stabilize the receptor, similar to constructs used in previous structural studies.^22, 24, 27^ In addition, a C-terminal StrepII purification tag and a N-terminal H-tag enhancing the translation initiation were attached to the receptor.^31^ The HFLβ1AR-G_s_ complex was formed by synthesizing HFLβ1AR in the presence of supplied G_s_, NDs (DOPG), and the agonist isoprenaline (Fig. S1). CF expression of HFLβ1AR in presence of all complex components resulted in higher HFLβ1AR-G_s_ complex formation efficiency compared to the formation achieved by incubating HFLβ1AR-NDs (DOPG) with the agonist and G_s_ after CF synthesis (data not shown). To further stabilize the complex, purified Nb35 was added in excess during CF synthesis and free nucleotides were hydrolyzed using apyrase before purifying the complex to homogeneity by IMAC and SEC.^32^ The resulting complex was imaged by cryo-EM to determine the structure of HFLβ1AR-G_s_ at an overall resolution of 3.3 Å (Fig. 1, Figs S2-S4 and Table S1).

**Fig. 1.**
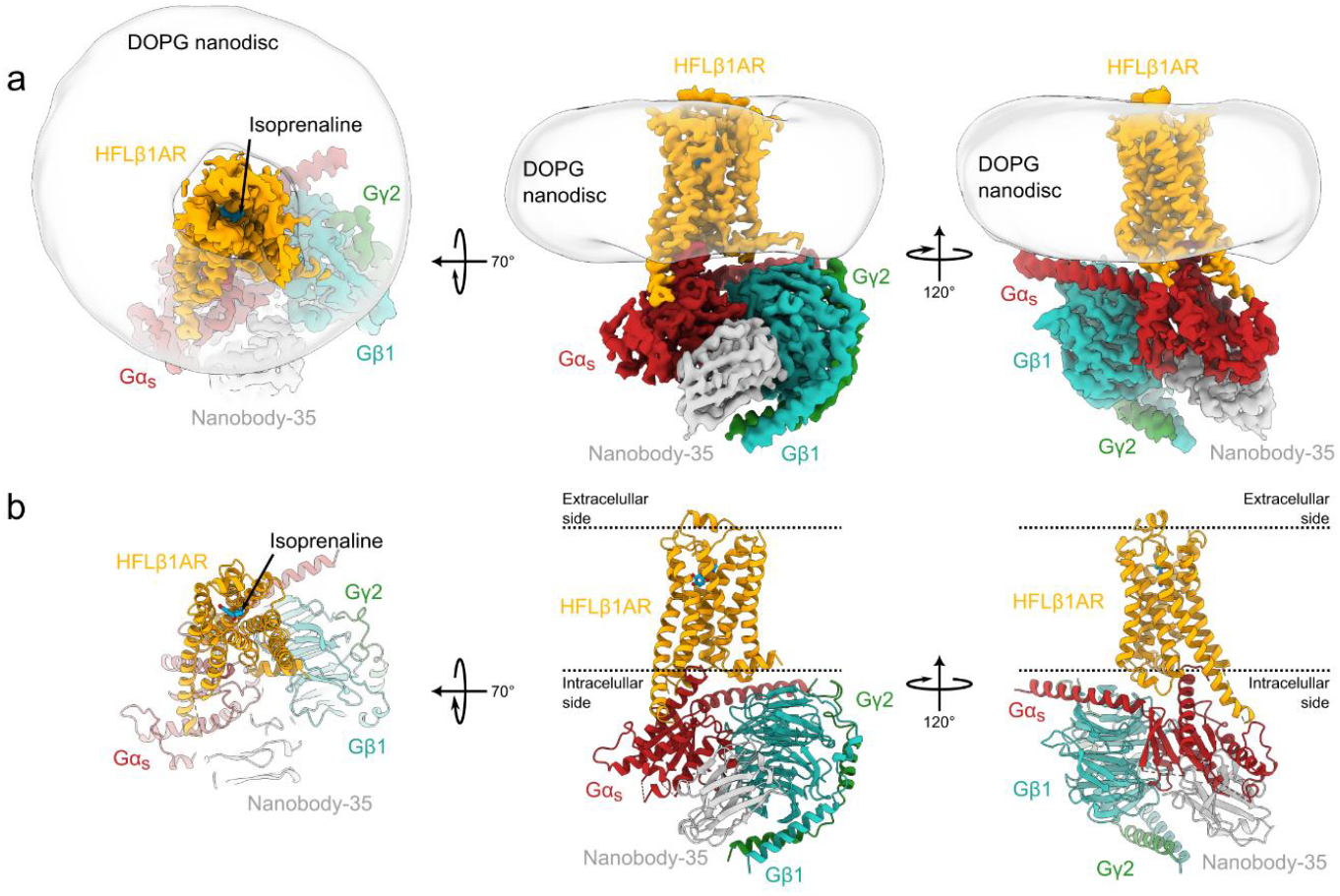
Cryo-EM structure of the isoprenaline-bound HFLβ1AR-G_s_ complex in ND. The ND density has been low-pass filtered and is shown as a transparent surface. The position of the agonist isoprenaline is indicated by an arrow. **a**. Cryo-EM density map of the complex. **b**. Ribbon model of the HFLβ1AR-G_s_ complex. Orange: HFLβ1AR; red: Gα_s_; cyan: Gβ; green: Gγ; grey: Nb35; blue: isoprenaline.

The density map shows uniform local resolution across the entire complex, with well-defined density at the interface between the receptor and the coupled G protein (Fig. S3). As observed in cryo-EM structures of other GPCR-G protein complexes, the loop region of the HFLβ1AR at the extracellular side of the transmembrane domain (TM) helix bundle exhibits the lowest resolution. In contrast, strong density is observed in the TM domains of the receptor, of the ligand isoprenaline within its binding pocket, and in the engaged G_s_ heterotrimer (Fig. S4), enabling us to build an atomic model of the isoprenaline-bound HFLβ1AR-G_s_ complex (Fig. 1). The receptor was modeled from residue Q55^1×29^ to the mutated palmitoylation site, C392A^8×59^, except for residues L266^ICL3^ to S312^ICL3^ of ICL3 and R313^6×24^ to R318^6×29^ of TM6, from which we observed no density. The heterotrimeric G_s_ protein and the stabilizing Nb35 were well resolved, except for regions of the Gα_s_ subunit known to exhibit conformational heterogeneity in the nucleotide free state.^33–37^ These regions include s1h1, H1, h1ha, the alpha helical domain (AHD), and hfs2 (residues A48^G.s1h1.2^-G206^G.hfs2.7^), s4h3 (residues S250^G.s4h3.1^-T263^G.s4h3.14^), HG (Q294^G.HG.1^-S306^G.HG.13^), and s6h5 (residues C365^G.s6h5.2^-A366^G.s6h5.3^) (the superscript numbers refer to the common Gα numbering (CGN) system).^36^

### Structural comparison of the HFLβ1AR-G_s_ complex with other active and inactive state β1AR structures

The structure of the CF-synthesized isoprenaline-bound HFLβ1AR-G_s_ complex closely resembles the previously published active-state structure of the N- and C-terminal as well as ICL3-truncated HΔβ1AR (PDBID 7BU6), bound to the endogenous agonist norepinephrine and stabilized by the G protein-mimetic nanobody Nb6B9 (root-mean-square deviation (RMSD) of 0.77 Å) (Fig. 2a).^10^ Like the HΔβ1AR, HFLβ1AR shows conformational changes in the TMs that are characteristic for class A GPCR activation when compared to the inactive structure of HΔβ1AR (PDBID 7BVQ) (Fig. 2b).^10, 38–40^ These include an 8.6 Å outward displacement of the intracellular end of TM 6, which is accompanied by more subtle inward movements of the intracellular parts of TMs 3, 5 and 7 to shape the intracellular transducer-binding pocket for the accommodation of the C-terminal α5 helix of the coupled G protein (Fig. 2b). These movements are associated with critical changes that are conserved in class A GPCR activation, including a slight shift of the W^6×48^ side chain in the C^6×47^W^6×48^xP^6×50^ motif, rearrangement of the P^5×50^I^3×40^F^6×44^ motif, movement of the N^7×49^P^7×50^xxY^7×53^ motif, and breakage of the ionic lock in the D^3×49^R^3×50^Y^3×51^ motif.^40–42^

**Fig. 2.**
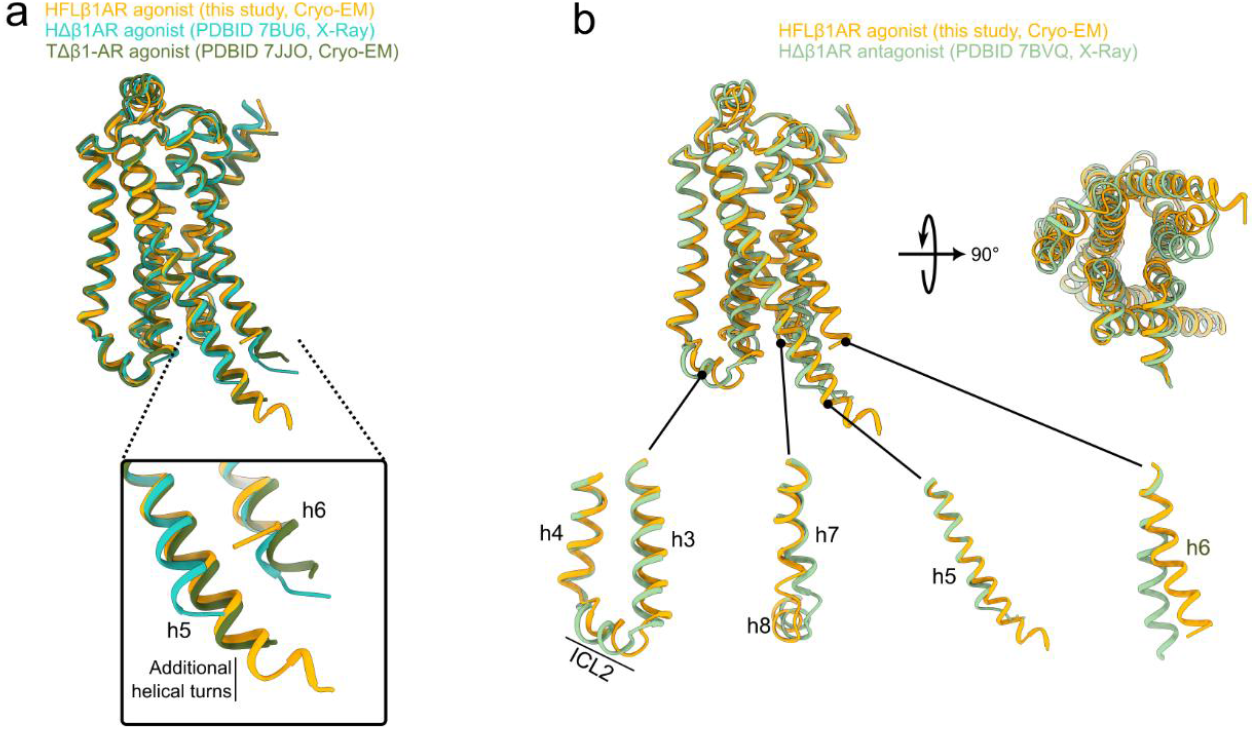
Comparison of active and inactive state structures of β1-adrenergic receptors. **a**. Superposition of HFLβ1AR with other β1 receptor structures in the active state. The extension of TM5 is highlighted in the panel. **b**. Superposition of HFLβ1AR with HΔβ1AR in the inactive state. The insets show the main conformational changes observed for each TM helix upon receptor activation.

Comparison of the HFLβ1AR-G_s_ complex with available structures of the ICL3-truncated turkey TΔβ1AR bound to G_s_ revealed a slightly different overall orientation of the G protein relative to the receptor (Fig. 3a, Fig. S5a). The N-terminal HN helix of the Gα_s_ subunit in the HFLβ1AR complex displays a rotation of ∼18° relative to the HN helix in TΔβ1AR-G_s_ (PDBID 8DCR). A similar rotation of Gβγ accompanies this change in Gα_s_ (Fig 3a, Fig. S5a). Despite differences in the relative orientation of the Gα_s_ subunit, the position of the C-terminal half of H5 within the receptor binding cavity remains strikingly similar across the different G protein-bound β1AR structures, allowing it to form critical interactions with the receptor core (Fig. 4). As seen in the TΔβ1AR-G_s_ complex structures, HFLβ1AR forms several polar interactions with H5 of Gα_s_ via residues in TMs 3, 5, and 6 (Fig. 4a).^24, 26, 43^ Specifically, the backbone carbonyl groups of I160^3×54^ and T161^3×55^ in TM3 engage in hydrogen bonding interactions with residues Q384^H5.16^ and R380^H5.12^ of H5 in Gα_s_, respectively. Additionally, K321^6×32^ and T325^6×36^ in TM6 establish ionic and H-bonding interactions with the C-terminal carboxyl group of L394^H5.26^ and the backbone carbonyl of E392^H5.24^ in the hook region of H5 (Fig. 4b). Furthermore, residues E250^5×64^, Q254^5×68^, and K257^5×71^ in TM5 contact H5 of Gα_s_ through H-bonding and ionic interactions with D381^H5.13^, Q384^H5.16^, and R385^H5.17^ (Fig. 4a). These polar contacts are consistent with previously published TΔβ1AR-G_s_ structures and mutagenesis data, which found that these core interactions are critical for β1AR-mediated G protein signaling.^24, 26, 43^

**Fig. 3.**
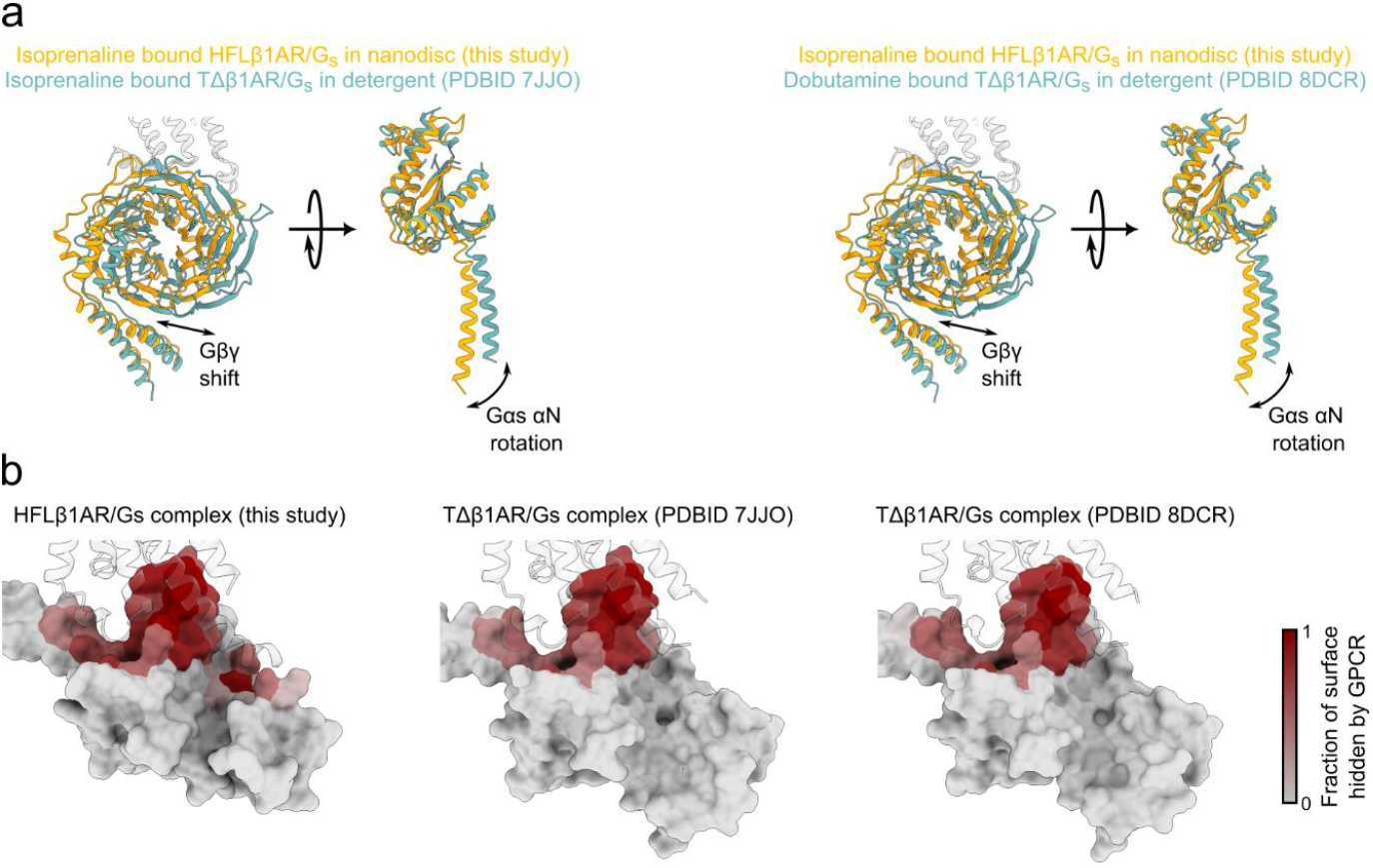
ICL3-dependent G protein orientations and receptor interaction interfaces between different β1AR/G_s_ complexes. **a**. Comparison of the orientation of the G protein relative to β1AR. The structures have been superimposed only on the receptor coordinates for all panels. For clarity, each panel shows either Gβγ or Gα_s_. **b**. Surface representation of Gα_s_, colored by the fraction of surface area hidden by binding to β1AR, which is shown as transparent ribbons.

**Fig. 4.**
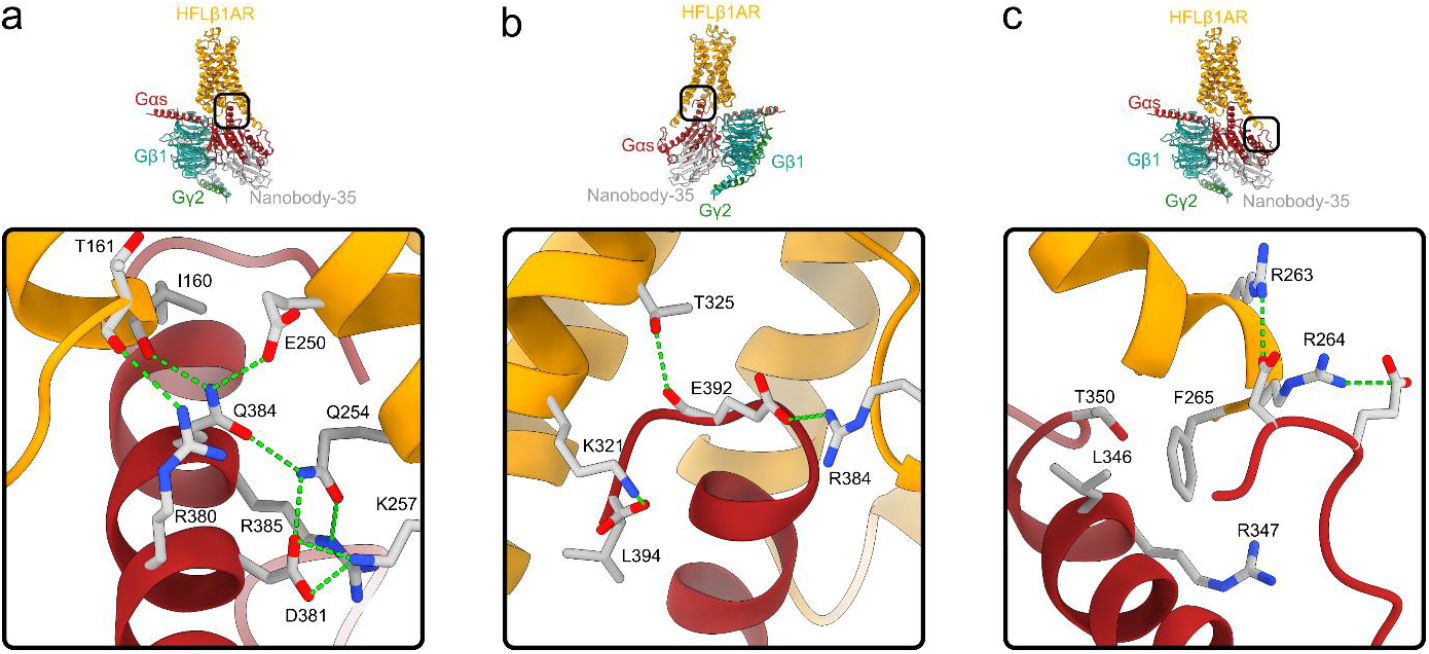
Main interface between HFLβ1AR and Gα_s_. **a, b**. Contacts between the α-helical (**a**) or hook (**b**) regions of H5 and HFLβ1AR. **c**. Contacts between the C-terminal end of TM5 and Gα_s_. Possible hydrogen bonding interactions between the two proteins are highlighted in green.

### HFLβ1AR ICL3 enhances the GPCR/G protein interface

The ICL3 of the turkey β1AR has been shown to modulate G protein coupling, thereby increasing G_s_ signaling and cAMP production.^28^ We therefore aimed to understand the structural basis of the involvement of ICL3 in G protein coupling. While the C-terminal region of ICL3 of HFLβ1AR could not be modeled due to the lack of density in the map, the N-terminal segment (C261^ICL3^-F265^ICL3^) was clearly resolved enabling detailed structural analysis of its interaction with the receptor-engaged G protein. In contrast to the active state structures of the ICL3-truncated derivatives HΔβ1AR bound to Nb6B9 and TΔβ1AR in complex with G_s_, the inclusion of ICL3 in our HFLβ1AR-G_s_ complex structure results in significant structural differences in the intracellular part of TM5. Compared to HΔβ1AR, the cytoplasmic end of TM5 in the HFLβ1AR structure is extended by nearly 3 helical turns, incorporating five residues of ICL3 (C261^ICL3^-F265^ICL3^) (Fig. 2a). For HΔβ1AR, which has a truncated ICL3 (C261^ICL3^ to L314^6×25^), TM5 is resolved only up to V255^5×69^, while residues K256^5×70^-S260^5×74^ are disordered. Additionally, the extended TM5 of HFLβ1AR adopts a slightly bent conformation, while the shorter TM5 in HΔβ1AR forms a straight helix (Fig. 2a). This bend in HFLβ1AR positions the C-terminal end of TM5 closer to TM6 (by approximately 5 Å, as measured between the Cα atoms of Q254^5×68^ in both structures). Notably, TM5 in the G_s_-coupled structure of TΔβ1AR (PDBID 7JJO), which features a similar truncated ICL3 (Y249^ICL3^-M283^6×28^) compared to HΔβ1AR, adopts a comparable bent conformation to that of the HFLβ1AR. However, it is only resolved up to D242^5×73^, making it six residues shorter than in the HFLβ1AR-G_s_ complex structure (Fig. 2a). As a result of this helical elongation and bending, TM5 in the HFLβ1AR-G_s_ complex facilitates more extensive interactions with the Gα_s_ subunit, specifically by binding into a cavity in the Ras domain of Gα_s_ formed by helices H4 and H5, as well as by the loops hgh4 and h4s6 (Fig. 3b). This leads to an increased interaction surface area between the receptor and Gα_s_ comprising 1427Å^2^ for HFLβ1AR-G_s_, compared to 1103Å^2^ for isoprenaline-bound TΔβ1AR-G_s_ (PDBID 7JJO), 1090Å^2^ for dobutamine-bound TΔβ1AR-G_s_ (PDBID 8DCR), or 1073Å^2^ for cyanopindolol-bound TΔβ1AR-G_s_ (PDBID 8DCS) (Fig. 3b).

Closer inspection of the interface revealed that the formation of the α-helical extension of TM5 creates additional interactions between residues in ICL3 and the G protein that are absent in the ICL3-truncated β1AR constructs. The guanidine groups of R263^ICL3^ and R264^ICL3^ in the HFLβ1AR-G_s_ structure are positioned near E322^hgh4.12^ and D323^hgh4.13^ (Fig. 4c, Fig. S5b) suggesting that they form ionic bonds with these residues in the hgh4 loop, which is not present in other G proteins. Additionally, F265^ICL3^, at the very end of the TM5 extension, packs against H4, forming apolar contacts with the side chains of L346^H4.16^, R347^H4.17^ and T350^h4s6.3^ (Fig. 4c) Notably, the consecutively residues R^ICL3^ and F^ICL3^ are highly conserved across β-adrenergic receptors, whereas the rest of the ICL3 region shows no or only low sequence similarity (Fig. S6). A comparison of our HFLβ1AR-G_s_ structure with available structures of the β2AR and β3AR, with intact ICL3s and coupled to G_s_, reveals that F^ICL3^ always forms similar hydrophobic contacts with residues of H4 (Fig. S7). In contrast, the side chain of the adjacent R^ICL3^ adopts different orientations but is still in the near vicinity of residues in the hgh5 loop or H4 of the Gα_s_. In the TΔβ1AR-G_s_ structure, these interactions are absent due to the truncation of ICL3, which only allowed structure determination of TM5 until position 5×73. However, despite the absence of ICL3, the core interactions between H5 and the TM bundle of the receptor are similar in both the HFLβ1AR-G_s_ and TΔβ1AR-G_s_ complexes. Given that the position of the G protein is different between the G_s_ complexes of HFLβ1AR and TΔβ1AR, while the core interactions are conserved, we speculate that the more extensive interface between the TM5-ICL3 region and the G protein results in the observed overall rotation of the G protein relative to the receptor. Indeed, previous work has shown that the presence of ICL3 results in different G protein coupling poses when compared with an ICL3-truncated receptor derivative.^28^

The different orientation of the G protein heterotrimer in the HFLβ1AR-G_s_ structure leads to an extended polar contact network between the receptor, Gα_s_, and the Gβ_1_ subunit (Fig. 4). This includes H-bond and ionic interactions between residues in H5 of Gα_s_ (D381^H5.13^, Q384^H5.16^ and E392^H5.24^) and TM5 (E250^5×64^, K257^5×71^) and H8 (R384^8×51^) of HFLβ1AR. Furthermore, H-bonds are formed between Q35^HN.52^ and R38^S1.03^ in the HN helix of Gα_s_ and the backbone carbonyl group of Q167^34×54^ in ICL2 of the receptor. Additionally, the rotation of the G protein brings residues R52 and D312 of Gβ_1_ in closer contact to ICL1 and H8 of HFLβ1AR, where they form an H-bond with Q90^12×51^ and an ionic bond with K385^8×52^ (Fig. 5), respectively. Since these contacts are absent in the TΔβ1AR-G_s_ structure, we speculate that while ICL3 is not essential for G protein coupling, it likely promotes a distinct G protein coupling pose. This pose facilitates additional interactions and a more extensive interface between the receptor and the G protein, which might further stabilize the β1AR-G_s_ complex resulting in enhanced cAMP production and downstream signaling. However, the structures of HΔβ1AR and TΔβ1AR have been solved in lipidic cubic phase and detergent, respectively, while the HFLβ1AR-G_s_ structure was obtained in ND (DOPG) environment. Therefore, we cannot rule out that these different membrane mimetics may contribute to the structural variations described above.

**Fig. 5.**
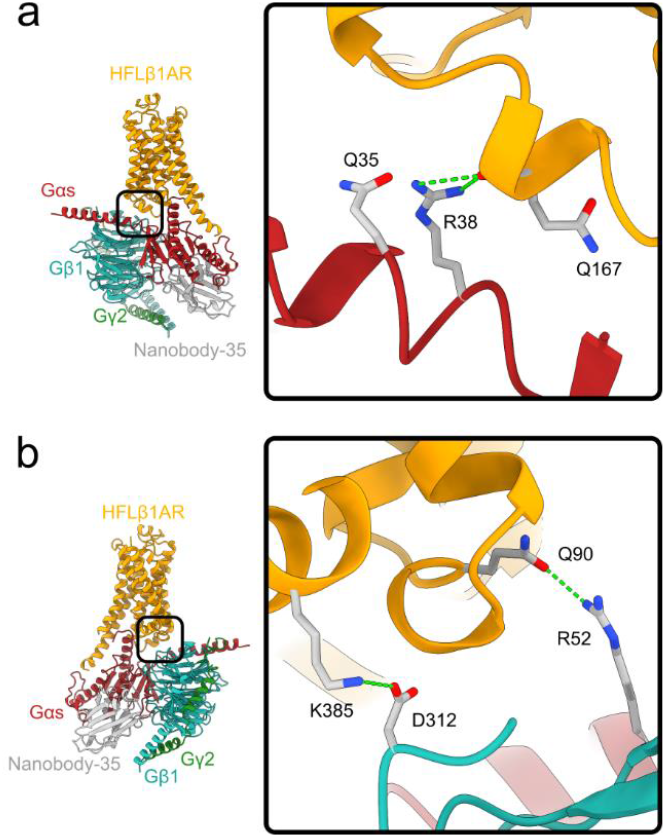
The presence of ICL3 leads to the formation of additional contacts between HFLβ1AR and the G protein. **a**. Contacts between the N-terminal helix of Gα_s_ and ICL2 of HFLβ1AR. **b**. Contacts between the ICL1 region of HFLβ1AR and Gβ. Possible hydrogen bonding interactions between the two proteins are highlighted in green.

### Mutational analysis of the extended HFLβ1AR-G_s_ protein-coupling interface

To directly test the relevance of the more extended interface between G_s_ and the full-length receptor HFLβ1AR for G protein signaling, we designed a derivative of wild-type Hβ1AR (Hβ1ARwt) similar to HΔβ1AR^10^ with residues C261^ICL3^-L314^6×25^ of ICL3 deleted and containing a C-terminus truncated at position 399 (= Hβ1AR^ΔICL3/CT^). Functional characterization using a cAMP signaling assay based on a cAMP bioluminescence resonance energy transfer (BRET) sensor (CAMYEL)^44^ revealed that these truncations reduced the EC_50_ value by approximately 130-fold, while showing a similar expression level compared to Hβ1ARwt or HFLβ1AR (Fig. 6, Fig. S8). This demonstrates that, while the ICL3 and the C-terminal regions are not essential for G protein coupling, they significantly modulate G protein activation. Despite the high sequence divergence between ICL3 of Hβ1AR and Tβ1AR and the potential additional effect of the C-terminus, our results align with previous findings on Tβ1AR, where truncation of ICL3 also impaired receptor-mediated G protein signaling.^28^

**Fig. 6.**
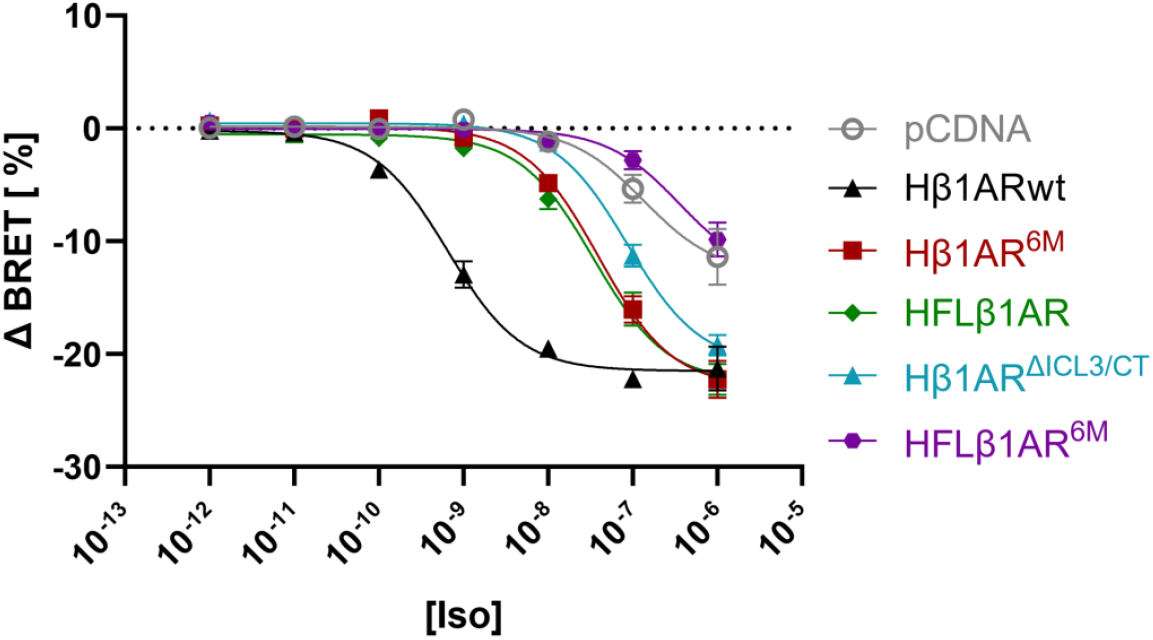
The presence of ICL3 and the additional receptor-G protein interactions enhance cAMP generation. Signaling assays for different constructs of the human β1AR. Signaling graphs represent the fit of grouped data ± SEM from three independent biological replicates. Iso, concentration of isoprenaline.

We next investigated the roles of polar and apolar interactions in the identified extended interface between Hβ1ARwt and the G_s_ heterotrimer in regulating G protein-coupling and activation. To this end, we generated single alanine mutations of residues in ICL3 (R263^ICL3^A, R264^ICL3^A, F265^ICL3^A) and TM5 (K257^5×71^A) that form polar contacts with the Gα_s_ subunit (Fig. S9a). In addition, we analyzed corresponding double (R263^ICL3^A/R264^ICL3^A) and triple (K257^5×71^A/R263^ICL3^A/R264^ICL3^A) mutants of the receptor (Fig. S9b). However, none of these mutations showed any significant impact on cAMP signaling (Fig. S9a, b). Notably, mutations E250^5×64^A in TM5 and Q167^34×54^A in ICL2 were not included in this study, as alanine substitutions of homologous residues in Tβ1AR have already been shown to significantly reduce the maximal G protein activation efficacy.^24, 26^ Mutations of Hβ1ARwt residues in H8 (K385^8×52^A) and in ICL1 (R88^12×49^A/Q90^12×51^A), which presumably interact with residues in the Gβ_1_ subunit decreased the potency by approximately 2- and 3-fold, respectively (Fig. S9c). However, the mutant Hβ1ARwt (R88^12×49^A/Q90^12×51^A) has a reduced surface expression level as well (Fig. S8). Next, we combined alanine substitutions in ICL1, ICL3 and H8 (R88^12×49^A/Q90^12×51^A/R263^ICL3^A/R264^ICL3^A/F265^ICL3^A/ K385^8×52^A) and tested their effect in both, Hβ1ARwt (Hβ1AR^6M^) and HFLβ1AR (HFLβ1AR^6M^) background (Fig. 6). Although these residues do not contribute to the core interactions between the receptor and the G protein found in previous structures of the β1AR, their alanine substitutions decreased the potency by 60-fold in the Hβ1ARwt background and 10-fold in the HFLβ1AR background. The smaller effect observed in the HFLβ1AR background is likely due to the 165-fold reduced potency of this construct compared to Hβ1ARwt, caused by the thermostabilizing mutations. In summary, these results implicate that the extended polar contact network in the HFLβ1AR-G_s_ structure, while not being essential for G protein coupling, modulate G protein coupling to the receptor and thus increase downstream cAMP signaling.

## DISCUSSION

Class A GPCR structures have provided key insights into their architecture and molecular mechanisms.^40, 45^ However, intrinsically disordered regions (IDRs) of GPCRs, such as segments in the N-terminus, C-terminal domain, and ICL3, are often truncated to enhance receptor expression and stability for structural studies. While their deletion generally preserves GPCR function, IDRs play crucial roles in modulating receptor signaling, desensitization, recycling, and targeting.^46, 47^ Serine and threonine residues in ICL3 and the C-terminus can be phosphorylated by GPCR kinases (GRKs) and other kinases like PKC and PKA, facilitating arrestin recruitment and receptor desensitization.^48, 49^ Additionally, both regions mediate interactions with GRKs and regulator of G protein signaling (RGS) proteins. Recent studies further highlight that the dynamic ICL3 and the C-terminus allosterically modulate receptor conformations to regulate receptor activity and G protein signaling.^50, 51^ Despite variations in length and primary sequence of ICL3 among GPCRs, it has been shown to autoregulate the activity of diverse receptors by sampling different conformational states that either occlude or expose the G protein-binding site, thus impacting G protein-coupling efficiency and specificity.^51–53^ Notably, GPCRs with an ICL3 longer than approximately 45 residues are proposed to exhibit greater G protein-coupling specificity than those with a shorter loop. For the Tβ1AR, hydrogen-deuterium exchange (HDX) mass spectrometry revealed that the receptor with an intact ICL3 couples more efficient to G_s_ and induces a distinct conformation of the nucleotide-free receptor-G protein complex compared to the truncated TΔβ1AR.^28^ However, the structural basis of the effect of ICL3 on the β1AR-G_s_ complex, which enhances cAMP production, remains unclear due to the absence of a full-length receptor structure in complex with its canonical G protein.

We previously employed direct cotranslational insertion of nascent GPCRs into preformed ND membranes by CF expression as suitable approach to determine the cryo-EM structure of the full-length human histamine 2 receptor (H_2_R) bound to the heterotrimeric G_s_.^19^ In the structure, ICL3 was fully resolved due to its short length of twelve amino acids connecting TMs 5 and 6. Here, we extend the application of the CF expression strategy to the HFLβ1AR, which possesses a longer ICL3 of 52 residues. Cryo-EM structure determination of the HFLβ1AR-G_s_ complex revealed similar GPCR-G protein core interactions compare to previously published structures of the ICL3-truncated turkey ortholog TΔβ1AR, involving the intracellular regions of TM3, TM5, TM6 and ICL2.^24, 26, 54^ While most of the ICL3 of HFLβ1AR is predominantly disordered in the structure due to its flexibility, the first five N-terminal residues of the loop are well structured, adopting an α-helical structure that extents TM5 of the receptor. Compared to available TΔβ1AR-G_s_ and HΔβ1AR-Nb6B9 complex structures, TM5 in HFLβ1AR is found to be 7-10 residues longer allowing it to protrude deeper into a pocket formed by H4 and H5, as well as the loops hgh4 and h4s6 of the Ras domain of Gα_s_.^10, 24, 26, 43^ This interaction is mainly found in G_s_-coupled receptor structures and has been proposed to determine the coupling selectivity of several class A GPCRs.^55^ In addition, this extended interface in the presence of ICL3 also likely causes the observed reorientation of the G protein relative to the receptor, enlarging the receptor-G_s_ interface compared to the reported Tβ1AR-G_s_ complexes.

The slight variation in receptor-G protein orientation enhances polar contacts between the receptor, the Gα_s_, and the Gβ_1_ subunit. This extended interaction network, facilitated by an intact ICL3, aligns well with previous findings showing increased coupling of an ICL3-containing Tβ1AR construct to miniG_s_ compared to the truncated derivative.^28^ Based on coarse-grained molecular dynamics simulations and hydrogen-deuterium exchange mass spectroscopy, Qiu et al. concluded that H5 of the miniG_s_ samples a different binding conformation to the surface of the receptor in the presence of ICL3. More specifically, a stronger H-bond interaction was predicted between Q237^5×68^ in TM5 of Tβ1AR and R385^H5.17^ in H5 of Gα_s_. This study further clarifies the stabilizing effect of ICL3 on the interaction of the HFLβ1AR-G_s_ complex by identifying new contacts between residues in TMs 5 and 8 with H5, ICL2 with HN, and ICL1 with H8 and the Gβ_1_ subunit. These findings provide new mechanistic insights into HFLβ1AR-G_s_ signaling that could inform the development of novel allosteric ligands, particularly ICL3-binding compounds that modulate the conformation of this loop region to influence G protein coupling and selectivity.^56^

Notably, the HFLβ1AR-G_s_ complex structure was obtained in DOPG NDs. Given that the lipid environment of GPCRs could influence ICL3 dynamics and stabilize distinct conformations, subtle modifications in the GPCR-G protein interface across different lipid or detergent environments are possible and cannot be ruled out.^57^ However, our recently published structure of the histamine H2 receptor in complex with G_s_ in a DOPG lipid environment closely resembles the receptor-G_s_ complex solved in detergent (PDBID 8YUT), with an RMSD of 1.12 Å, suggesting that differences in the environment may have a relatively minor impact on the structure of GPCR-G protein complexes.^19, 58^

The disordered terminal domains of GPCRs play crucial roles in ligand binding, as well as G protein and arrestin coupling.^50, 59–64^ Although these domains were not resolved in the HFLβ1AR structure, their inclusion in the structure of other class A receptors have provided novel structural insights into their functional role. Ordered packing of the N-terminus at the extracellular side has been observed in structures of a number of GPCRs such as the cannabinoid receptor 1 (CB1), the µ-opioid receptor (µOR), the angiotensin II receptor type 1 (AT1R), the lysophosphatidic acid receptor 1 (LPA1), the chemokine receptor CXCR4, and the orexin 1 receptor (OX1R),^65–70^ where the N-terminus contributes to ligand binding or recruitment. Additionally, in the cryo-EM structure of the muscarinic M1 receptor (M1R) and rhodopsin in complex with their canonical G proteins, a segment of the C-terminus binds to a cleft between the Gα and Gβ subunit of the engaged heterotrimeric G protein, suggesting that it plays a role in G protein coupling and specificity.^64, 63^ Notably, in the HFLβ1AR-G_s_ complex, close inspection of the cryo-EM density at higher contour levels suggests that H8 forms an extended α-helix, appearing 2-3 helical turns longer compared to previous β1AR structures in complex with G_s_ or Nb6B9 (Fig. S4c).^10, 24, 26, 43^ Although this extension was not included in the final model due to the weak density presumably caused by the increased dynamics of this region in the absence of H8 palmitoylation, it highlights how the inclusion of the intrinsic dynamic domains can uncover notable structural features. The extension may result from ionic interactions between basic residues in the proximal C-terminal region of H8 (R395-H402) and phospholipid head groups of the ND, though its functional significance remains unclear. Since cell-based expression systems often require truncation of disordered termini or loops to enhance receptor expression and biochemical stability for structural characterization, the CF expression system based on *E. coli* lysates, used in this study, offers a complementary approach, enabling the rapid production of high quality, full-length GPCR samples for structure determination.

## RESOURCE AVAILABILITY

### Lead contact

Further information can be obtained by the lead contact, Frank Bernhard.

### Materials availability

This study did not generate new unique reagents.

## Data and code availability

The EM map for the complete HFLβ1AR molecule has been deposited in the EMDB under accession code EMD-19683. Atomic coordinates for HFLβ1AR have been deposited in the Protein Data Bank under the accession code <u>PDB</u> 8S2T. <u>Source data</u> are provided with this paper.

## ACKNOWLEDGEMENTS

We thank Birgit Schäfer and Alexander Strubel for helpful discussions and technical assistance. We are further grateful to Betsy White and Hannes Schihada for help with G protein expression and the cAMP signaling assay, respective. The work was funded by the DFG project BE1911/9-1, by the Center for Biomolecular Magnetic Resonance and by the LOEWE project GLUE LOEWE/2/12/519/03/05.001(0014)/71 of the state of Hessen. LOEWE GLUE financed the theses of ZK.

## AUTHOR CONTRIBUTIONS

Sample preparations and biochemical studies were done by ZK, PD, AG and DH. FM, UE, ZK, AS and ASF performed cryo-EM data acquisition and analysis. DH provided heteromeric G-proteins. PD prepared and characterized mutants using the cAMP assay under supervision of DH. VD and JK provided essential support and infrastructure. FB and DH conceived the project. All authors contributed to manuscript writing, data analysis, reading and approving the final version of the manuscript.

## DECLARATION OF INTERESTS

FM and JK are shareholders of Cube Biotech, which sells products and services related to membrane protein characterization. All other authors declare no competing financial interests. The article is the authors’ original work, has not received prior publication and is not under consideration for publication elsewhere.

## SUPPLEMENTAL INFORMATION

*Figures S1–S7 and Table S1*

## METHODS

### *E. coli* lysate preparation

*E. coli* S30 lysates were prepared following established protocols.^71, 72^ In summary, 200 mL LB media was inoculated with a single colony of *E. coli* A19 cells and incubated overnight at 37

°C with shaking at 180 rpm. A 10 L fermenter containing YPTG medium (16 g/L peptone, 10 g/L yeast, 5 g/L NaCl, 100 mM glucose, 22 mM KH_2_PO_4_, 40 mM K_2_HPO_4_) was inoculated with 100 ml of the pre-culture and approx. 1 mL antifoam Y-30 emulsion (Sigma-Aldrich, Taufkirchen, Germany). The cells were grown at 37 °C with frequent stirring at 300 rpm until the OD600 reached 3.5-4. Following rapid cooling to 20 °C, the culture was harvested by centrifugation at 5,000 × g and 4 °C for 30 min. The cells were resuspended in 300 mL S30 buffer A (14 mM Mg(OAc)_2_, 60 mM KCl, 6 mM β-mercaptoethanol, 10 mM Tris-acetate, pH 8.2) and subjected to centrifugation at 10,000 × g and 4 °C for 10 min. This washing process was repeated twice, with the final centrifugation extended by 30 min. The pellet was resuspended in 110 % (w/v) S30 buffer B (14 mM Mg(OAc)_2_, 60 mM KCl, 1 mM DTT, 10 mM Tris-acetate, pH 8.2) and the cells were lysed using a French press at a constant pressure of 20,000 psi. The disrupted cells were centrifuged twice at 30,000 × g and 4 °C for 30 min. The resulting supernatant was adjusted to 400 mM NaCl and incubated at 42 °C for 45 min to allow for mRNA run-off. Subsequently, the lysate was dialyzed against 5 L S30 buffer C (14 mM Mg(OAc)_2_, 60 mM KAc, 0.5 mM DTT, 10 mM Tris-acetate, pH 8.2) for 3 h followed by another dialysis for 16 h at 4 °C. A final centrifugation step was performed at 30,000 × g and 4 °C for 30min. The supernatant was collected, aliquoted, flash frozen in liquid nitrogen and stored at −80 °C.

### Production of MSP1E3D1

MSP1E3D1 was synthesized in *E. coli* T7 express cells (New England Biolabs, Frankfurt, Germany) and purified using its N-terminal His-tag.^73^ In order to remove the N-terminal His_6_-tag, the protein underwent TEV digestion, followed by reverse IMAC. The MSP1E3D1 solution was adjusted to 1 mM DTT, and TEV protease was added at an MSP1E3D1 to TEV ratio of 1:25. The mixture was subjected to overnight dialysis at 4 °C in cleavage buffer (1 mM DTT, 0.5 mM EDTA and 50 mM Tris-HCl, pH 8.0). Prior to loading onto a pre-equilibrated HiTrap™ IMAC FF column (Cytiva, Munich, Germany), the mixture was centrifuged at 20,000 x g for 10 min at 4 °C. The flow-through was collected, and the column was washed with 10 CV of equilibration buffer (20 mM imidazole, 100 mM NaCl, 20 mM Tris-HCl, pH 8.0). The flow-through and wash fractions were pooled and concentrated to a final concentration of 3-5 mg/mL using Amicon ultrafiltration units (10 kDa MWCO, Merck Millipore, Darmstadt, Germany). The protein solution was dialyzed twice overnight at 4 °C against 5 l of buffer containing 10 % (v/v) glycerol, 300 mM NaCl, and 40 mM Tris-HCl, pH 8.0. The protein was flash-frozen in liquid nitrogen and stored at – 80 °C until further use.

### Nanodisc formation

NDs were prepared as described previously. Briefly, purified MSP1E3D1 was incubated with the corresponding lipid and supplemented with 0.1 % DPC. NDs were assembled at specific MSP1E3D1 to lipid ratios, depending on the lipid used: 1:80 for DOPG.^73, 74^ The solutions were incubated at room temperature (RT) for 1 h with gentle stirring, then dialyzed three times for at least 12 h at RT against a buffer containing 100 mM NaCl, 10 mM Tris-HCl, pH 8.0. To remove potential precipitates after dialysis, the mixture was centrifuged at 20,00 x g at 4 °C for 20 min. NDs were concentrated to 500 – 1000 µM using Centriprep concentrating units (10 kDa MWCO, Merck Millipore, Darmstadt). Lipids and DPC were purchased from Avanti Polar Lipids (Alabaster, USA).

### Cell-free expression

The expression of CF protein was conducted in lysates of the *E. coli* strain A19 using a two-compartment set-up in accordance with the methodology previously described in detail.^31, 75^ Analytical-scale reactions were conducted in 24-well plates with Mini-CECF reactors. For preparative-scale reactions, 3 mL Slide-A-Lyzer devices (10 kDa MWCO, Merck Millipore, Darmstadt, Germany) were combined with custom-made Maxi-CECF containers. The reaction mixture to feeding mixture ratios were 1:15 for analytical and 1:17 for preparative scale reactions. The reaction was performed in a previously determined Mg^2+^ optimum (16–20 mM Mg(OAc)_2_), with additional components including 1 mM of each amino acid, 20 mM acetylphosphate, 20 mM phosphoenolpyruvate, 0.1mg/ mL folinic acid, 1× complete protease inhibitor (Roche, Penzberg, Germnay), 270 mM KOAc, 3 mM GSH, 1 mM GSSG, 1.2 mM ATP, 0.8 mM each CTP, GTP, UTP, and 100 mM HEPES-KOH pH 8.0. The reaction mixture (RM) also included the *E. coli* S30 lysate, 15 ng/μL DNA template, 0.3 U/μL RiboLock RNase inhibitor (ThermoScientific, Langenselbold, Germany), 10–20 U T7 RNA polymerase, 0.04 mg/mL pyruvate kinase and 0.5 mg/mL *E. coli* tRNA (Roche, Penzberg, Germany). All constructs were co-translationally inserted into preformed NDs, which were provided to the RM. The reactions were incubated at 30 °C for 16 to 20 h with gentle shaking. After expression, the RMs were harvested by centrifugation at 18,000 x g at 4 °C for 10 min to remove any precipitates.

The purification of GPCR/ND complexes were performed via StrepII-Tactin affinity chromatography using a pre-equilibrated gravity flow column in buffer A (100 mM NaCl, 20 mM HEPES, pH 7.4). RMs were diluted in a 1:3 ratio in buffer A prior to loading on the columns. The columns were washed with 10 CV buffer A and eluted in 4 CV buffer A + 25 mM d-desthiobiotin. Samples were concentrated in Amicon Ultra 0.5 mL units (50 kDa MWCO, Merck Millipore, Darmstadt, Germany).

### Size exclusion chromatography (SEC)

SEC analysis was performed at 16 °C using an ÄKTA purifier system (Cytiva, Munich, Germany) and Increase Superose 6 5/150 or 3.2/300 columns (Cytiva, Munich, Germany). Prior to loading with the affinity-purified protein samples, the columns were equilibrated with sterile-filtered, degassed and pre-cooled buffer A. The flow rate was set to 0.15 mL/min for the Superose 6 5/150 column and 0.05 mL/min for the Superose 6 3.2/300 column. UV absorbance was measured at 280 nm. Data were plotted using GraphPad Prism (v 10).

### Preparation of Nb35

Nb35 was expressed and purified following standard protocols.^32^ In short, Nb35 was expressed in *E*.*coli* BL21 cells. Following cell lysis, it was purified using IMAC. SEC was carried out on a Superdex 200 10/300 gel filtration column (GE Healthcare) in 150 mM NaCl, 20 mM HEPES pH 7.5. Purified Nb35 was concentrated, flash-frozen and stored at -80 °C until further use.

### Cryo-EM sample preparation

The HFLβ1AR-G_s_ complex was formed co-translationally in the CF reaction, as described previously.^19^ For this purpose, the reaction was supplemented with the final concentrations of the following: 15 ng/µL HFLβ1AR DNA template, 70 µM NDs (DOPG) without His-tag, 10 µM purified G_s_ heterotrimer, 15 µM Nb35 and 0.5 mM isoprenaline. The reactions were incubated at 30 °C for 16 h with gentle shaking. Following incubation, samples were subjected to centrifugation at 18,000 x g for 10 min at 4 °C. The sample was incubated on ice with 1 U/µL apyrase for 90 min. The HFLβ1AR-G_s_ complex was then purified via IMAC at 4 °C. The samples were diluted 1:3 in IMAC buffer A (100 mM NaCl, 20 mM HEPES, pH 7.4). They were subsequently loaded onto a pre-equilibrated gravity-flow IMAC column, and reloaded twice. The column was then washed with 4 CV each of IMAC buffer A and IMAC buffer B (IMAC buffer A + 30 mM imidazole). The complexes were eluted in 4 CV IMAC elution buffer (IMAC buffer A + 300 mM imidazole) and samples were subjected to concentration using Amicon Ultra 0.5 mL units (50 kDa MWCO, Merck Millipore, Darmstadt, Germany). Subsequently, SEC was conducted as described above using the Superose 6 5/150 column. Fractions containing the complex were then pooled and concentrated using Amicon Ultra 0.5 mL units (50 kDa MWCO, Merck Millipore, Darmstadt, Germany).

### Cryo-EM grid freezing and data collection

Freshly purified HFLβ1AR-G_s_ complex was used for cryo-EM analysis. Sample vitrification was carried out using a Vitrobot (FEI) operated at 100% humidity and 13°C. A 3.5 µL sample concentrated to 0.8 mg/ml was placed onto a glow-discharged UltraAU foil R1.2/1.3 300 mesh (Quantifoil) grid. The excess sample was blotted away for 10 s with -5 force and the grid was immediately plunged into liquid ethane. The grids were then transferred into a Titan Krios microscope equipped with an XFEG operating at 300 kV, a K2 summit direct detector (Gatan), and a GIF Quantum energy filter (Gatan). All images were collected at 165,000x magnification corresponding to a pixel size of 0.81 Å, defocus values between -1.0 and -2.5 µm, and a 20-eV slit width for the energy filter. Exposures lasted 9 s, resulting in a total dose of 65 e^-^/Å^2^. For each exposure, a 45-frame movie was collected with the camera operated in counting mode. Data collection was handled by SerialEM.^76^

### Cryo-EM data processing and model building

We collected a total of 1,654 movies, which were aligned and dose corrected using Relion’s MotionCor implementation.^77^ After CTF estimation with CTFFIND4,^78^ we discarded the images with bad ice or bad CTF estimation, resulting in 1,377 images selected for further processing. We used Topaz or particle selection, and Relion 4 for classification and refinement.^79, 80^ Initially, we selected 330,942 particles using the general model provided by Topaz. After several rounds of 2D classification and 3D refinement used for recentring, we selected a subset of 5,867 clean particles from 100 images, which we used to train an *ad hoc* model for our data set. With the trained model, we selected a total of 235,480 particles, which were extracted into 384px boxes binned down to 192px (1.62 Å/px). Several rounds of 2D classification combined with 3D refinement for recentring resulted in a subset of 124,176 particles. 3D refinement with this subset, using sidesplitter for regularization, resulted in a 4.0-Å-resolution reconstruction of the complex.^81^ We then subtracted the signal coming from the ND^54^ and ran one round of 3D classification with 3 classes and fast subsets, resulting in a single class with clear GPCR density containing 74,266 particles. After magnification and per-micrograph CTF correction,^82^ refinement with these particles resulted in a 3.7-Å-resolution reconstruction of the complex. We used this reconstruction for Bayesian polishing,^77^ after which we re-extracted the particles binned this time to 300 px (1.036 Å/px). A new round of ND subtraction and per-particle CTF refinement resulted in a 3D reconstruction of 3.5-Å-resolution. A final round of refinement using the original polished particles and blush regularization in Relion 5 produced a 3.3-Å-resolution map.^83^ Fig. S2 shows a schematic view of the data processing strategy. Local resolution was estimated with mFSC within SPHIRE.^84, 85^ Directional resolution was estimated using the 3D-FSC package.^86^

For the initial model of the complex, we used ChimeraX to rigid body fit G_s_ and Nb35 from PDBID 6E3Y and a colabfold model of HFLβ1AR into the map.^87–89^ We refined this model by iterative rounds of manual adjustment in Coot and the iterative-density-guided rebuilding protocol implemented in Rosetta.^90, 91^

### β1AR functional test with CAMYEL biosensor

A codon optimized sequence of the human β1AR with N-terminal HA- and FLAG-tag was ordered from Twist Bioscience and subcloned into pCDNA vector using EcoRI and XbaI restriction sites. Mutations were introduced with the Quick-Change approach with primer including one or multiple mutations and amplification using PrimeStar MAX DNA polymerase (Takara Bioscience). HEK 293A cells were grown in Dulbecco’s modified Eagle’s medium (DMEM) supplemented with 10% fetal bovine serum, 100 U/ml penicillin/streptomycin, 2 mM L-glutamine at 37 °C in 5% CO_2_. All transfections were carried out as follows: 48 h before the experiments, cells were transfected in suspension using polyethylenimine at 3:1 PEI/DNA ratio. 500 ng DNA per receptor construct and 200 ng CAMYEL biosensor were diluted into OptiMEM and adjusted to 1µg/ ml cells with empty pCDNA vector.^44^ For BRET experiments (∼3×10^4 cells/well) were seeded in white 96-well-plates for ELISA assays (∼6×10^4 cells/well) were seeded to PDL coated transparent flat bottom 96-well-plates.

48 h after transfection cells were washed 2 times with Hank’s balanced salt solution, pH 7.4. To measure concentration-response curves, luciferase substrate coelenterazine 400a (5 µM, Nanolight Technology) and ICI 118,551 (100 nM; Tocris Bioscience), as selective β2AR inhibitor, were added to the cells and incubated for 5 min at 37 °C. BRET was measured using a CLARIOStar Microplate reader using monochromator (acceptor 524.9-70 nm and donor 400-80 nm) at 37 °C. After 3 initial reads, indicated concentration of isoprenaline (Sigma-Aldrich) were added and BRET decrease was detected for 15 more reads. Absolute BRET ratio was calculated by dividing detected emission form the acceptor through detected emission from the donor channel. The average of three consecutive BRET values before ligand stimulation were defined as basal BRET. Afterwards raw ΔBRET was calculated for each well after ligand addition as a percent over basal ([(Ratio_stim_ − Ratio_basal_)/Ratio_basal_] × 100). From this, the average raw ΔBRET of wells where only vehicle was added, was subtracted for final ΔBRET.

For quantification of cell surface receptor expression, after 48 h HEK293A cells transfected with pCDNA or N-terminally HA-tagged β1AR receptor constructs were incubated with 50 µL of M1 anti–HA-tag antibody (1 µg/mL,) in 1% BSA–PBS for 1 hour at 4°C. Afterwards, five washing steps with 0.5% BSA–PBS followed. Cells were incubated then with an HRP–conjugated antibody (92 ng/ml; Cell Signaling Technologies) in 1% BSA–PBS for 1 hour at 4 °C. The cells were washed again five times with 0.5% BSA–PBS, and 100 µL of the peroxidase substrate 3,3′,5,5′-tetramethylbenzidine was added. Subsequently, the cells were incubated for 20 min at room temperature and 100 µL of 2 M HCl was added. The absorbance was read at 450 nm using a BMG ClarioStar plate reader.

