## Supplemental figures S1-S9 and Table S1 for "Cryo-EM structure of a cell-free synthesized full-length human β1-adrenergic receptor in complex with G_s_"

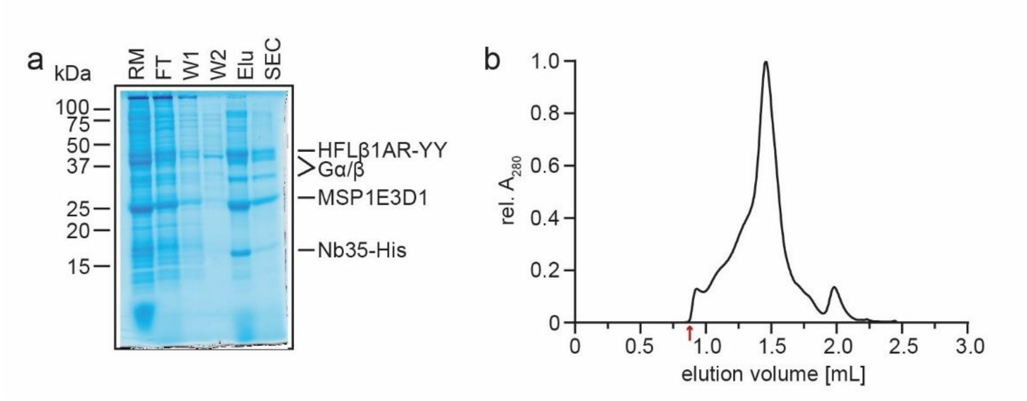

**Fig. S1. Purification of the HFLβ1AR-$G_s$ complex in nanodiscs. a.** SDS-PAGE of IMAC-purified HFLβ1AR-$G_s$ complex samples in NDs (DOPG) prepared after CF synthesis in presence of isoprenaline. The gel was stained with colloidal Coomassie-Blue staining solution. RM: reaction mix, FT: flow-through, W1: first wash, W2: second wash, Elu: elution, SEC: sample of the main peak after SEC. **b.** SEC profile of IMAC purified HFLβ1AR-$G_s$ complex in NDs (DOPG) complex. Gel filtration was carried out on a Superose 6 3.2/300 column. The column's void volume is indicated by an arrow.

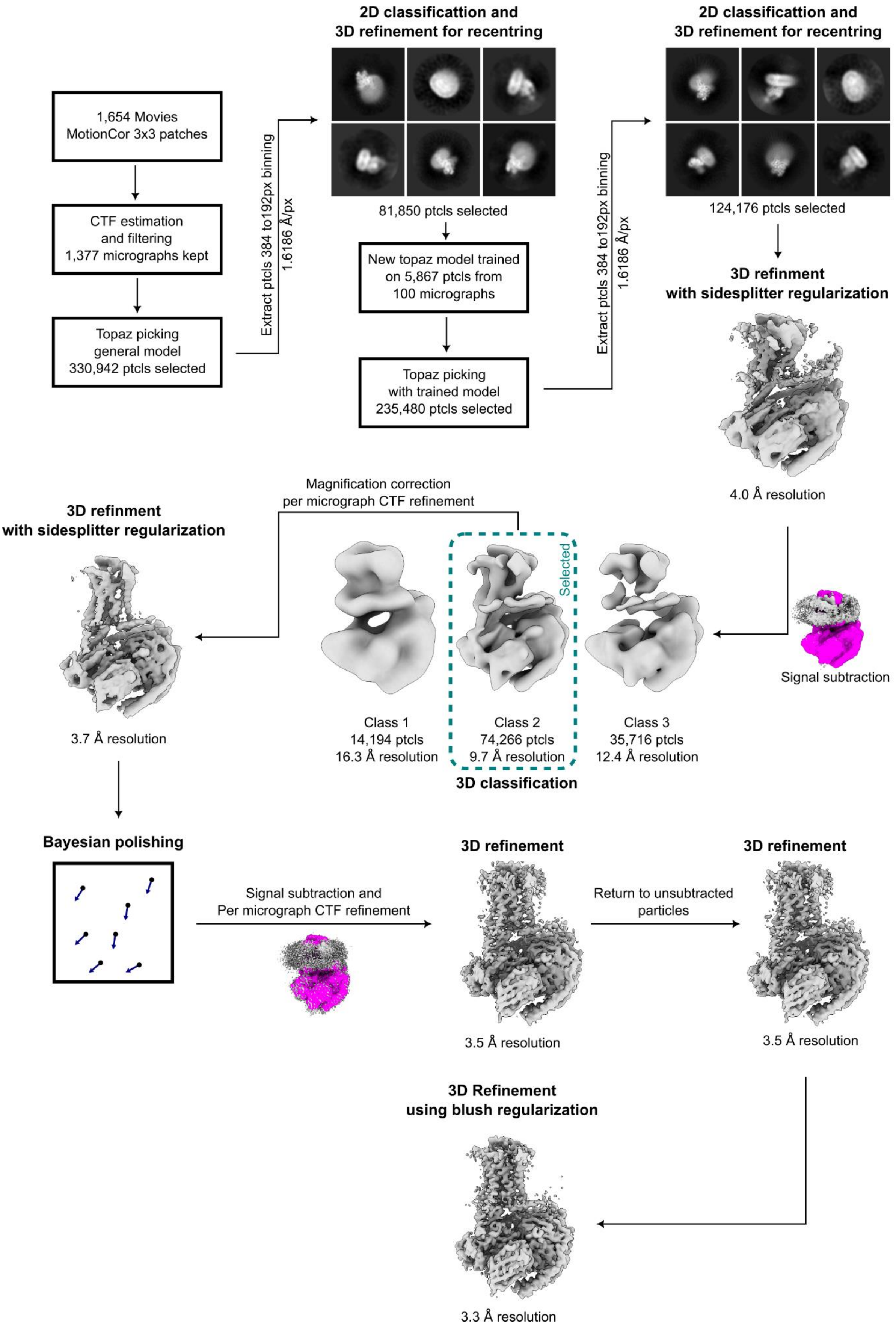

**Fig. S2. Cryo-EM data processing workflow**.

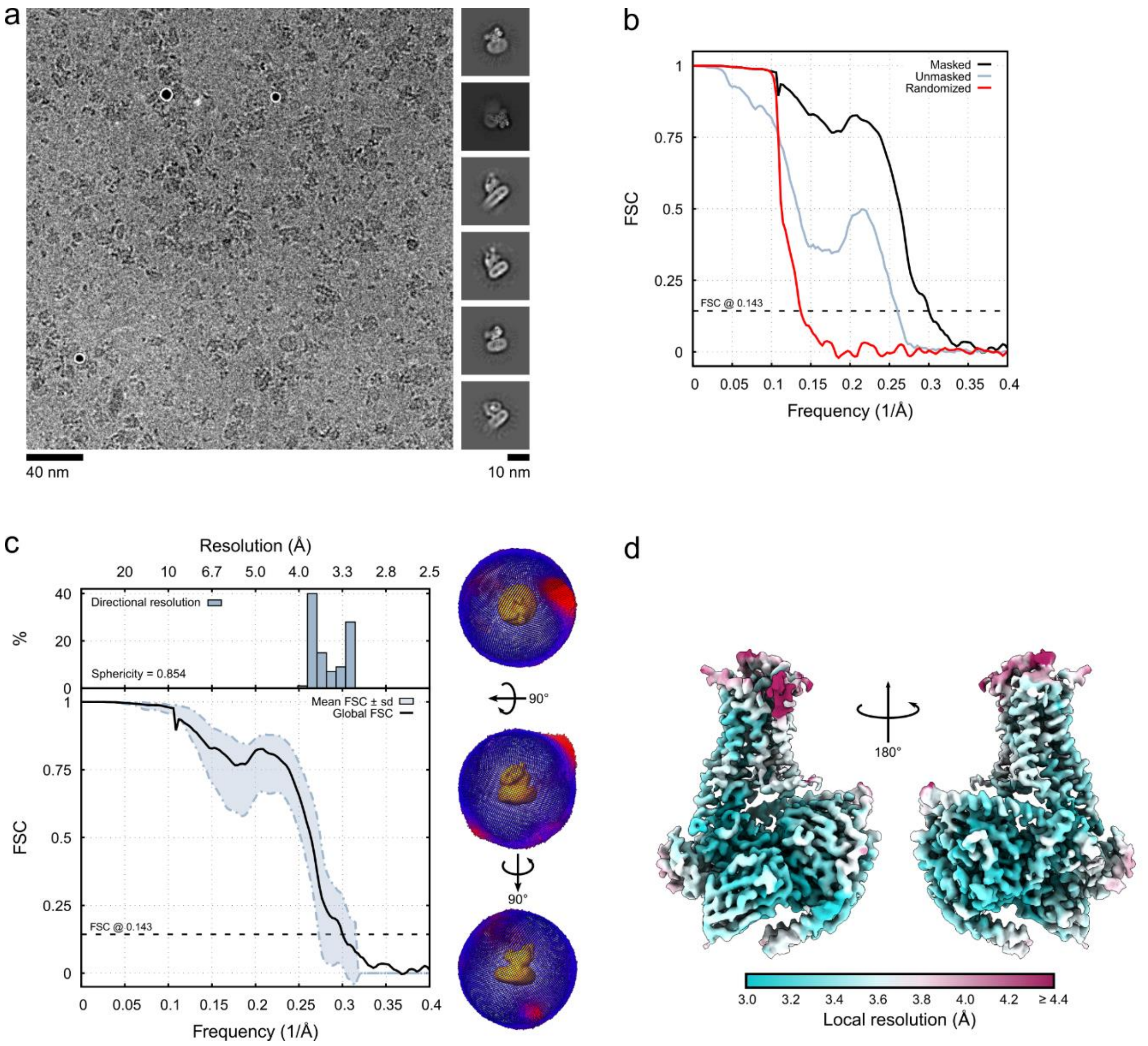

**Fig. S3. Resolution of the HFLβ1AR-$G_s$ complex in NDs**. **a.** Representative cryo-EM micrograph and selected 2D-class averages. **b and c**. Global (**b**) and directional (**c**) FSC curves for the reconstruction. **d.** 3D reconstruction of the HFLβ1AR-$G_s$ complex colored according to the local resolution.

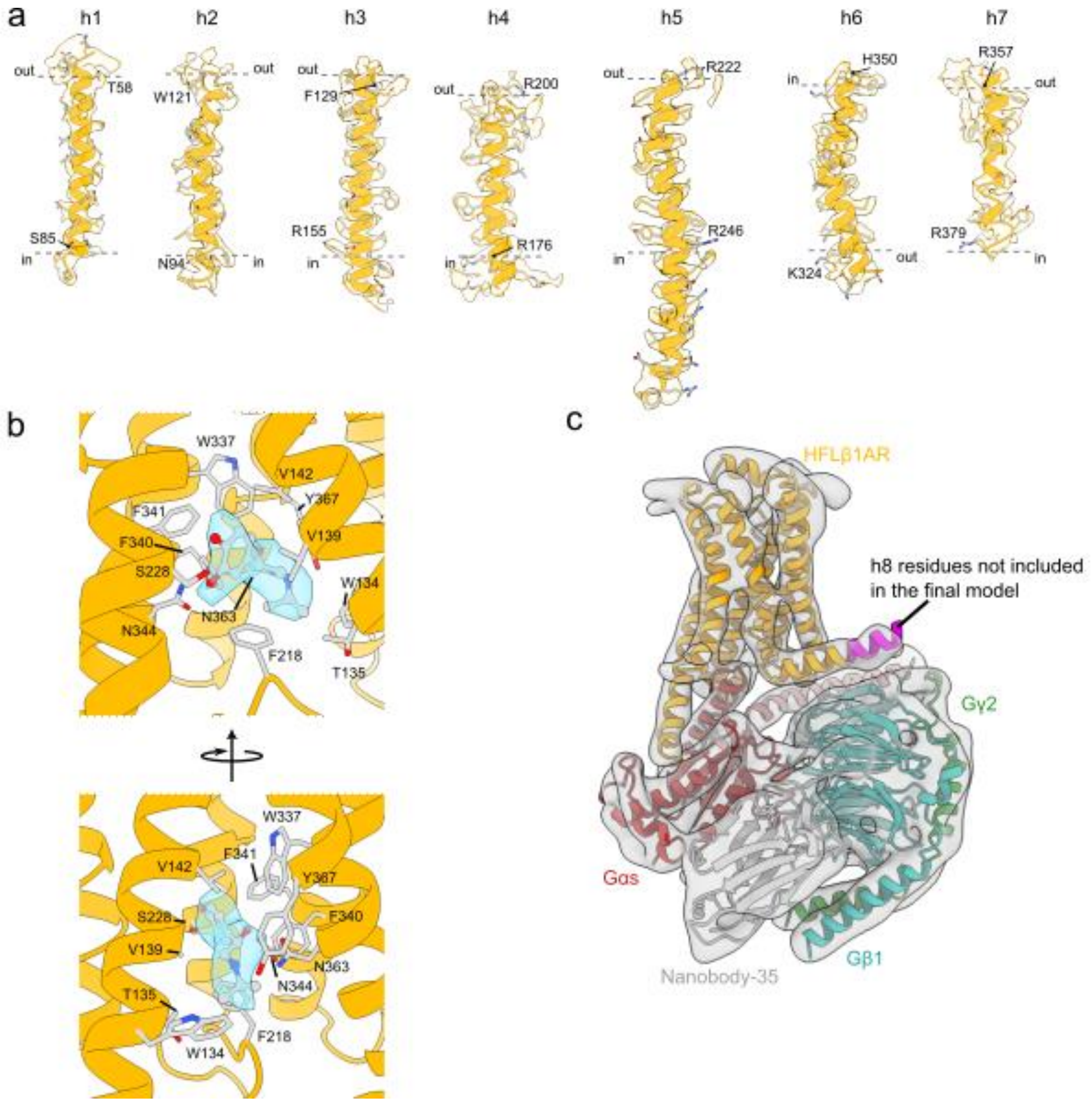

**Fig. S4. Cryo-EM densities of the HFLβ1AR-G$_s$ complex**. **a.** Densities of the seven transmembrane helices of HFLβ1AR. **b.** Densities of isoprenaline in the ligand binding pocket. **c.** Density of the complex filtered to 8 Å, showing additional density for helix 8 (H8) of HFLβ1AR.

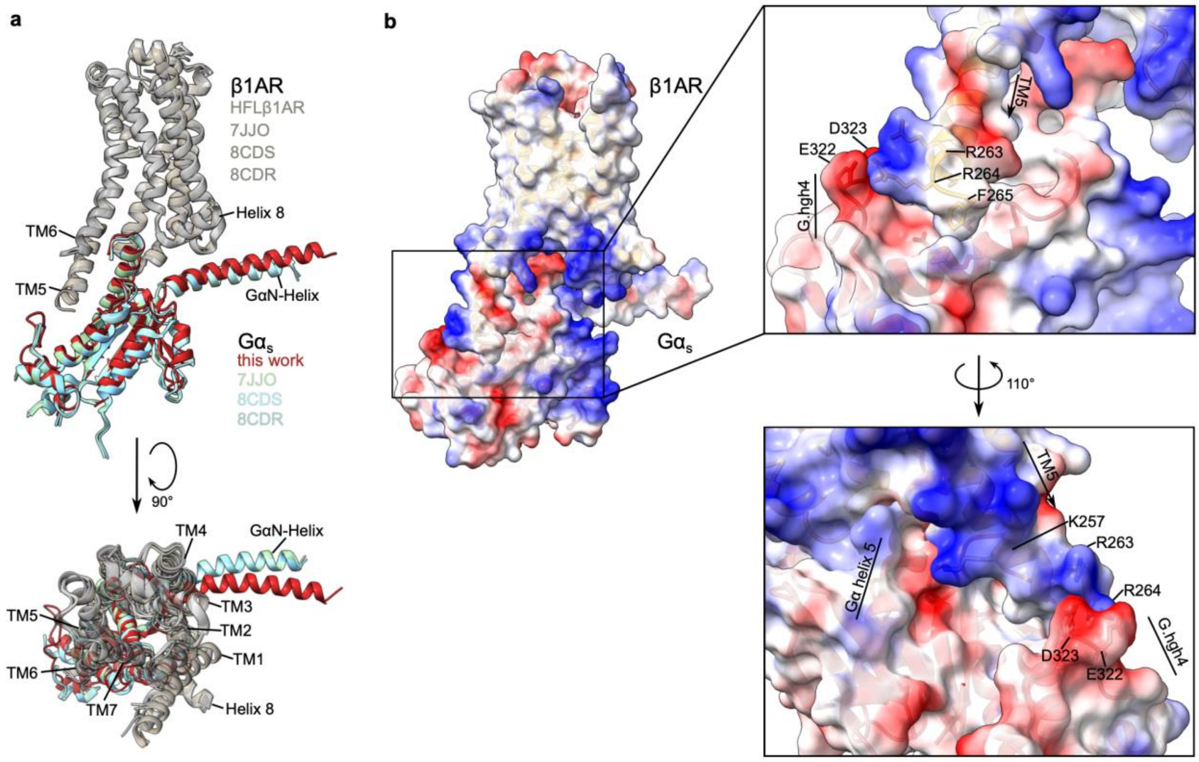

**Fig. S5. ICL3-induced changes in the receptor-G protein interface in the HFLβ1AR-$G_s$ structure. a.** Structural comparison of the relative G protein orientation in the full-length human HFLβ1AR-$G_s$ complex and truncated TΔβ1AR-$G_s$ complexes. **b**. Representation of the electrostatic surface area of HFLβ1AR and the engaged $G\alpha_s$ subunit. The extended cytoplasmatic end of TM5 of the receptor shows a number of positively charged residues (blue) that form electrostatic interactions with negatively charged residues in the G.hgh4 loop of $G\alpha_s$ (red).

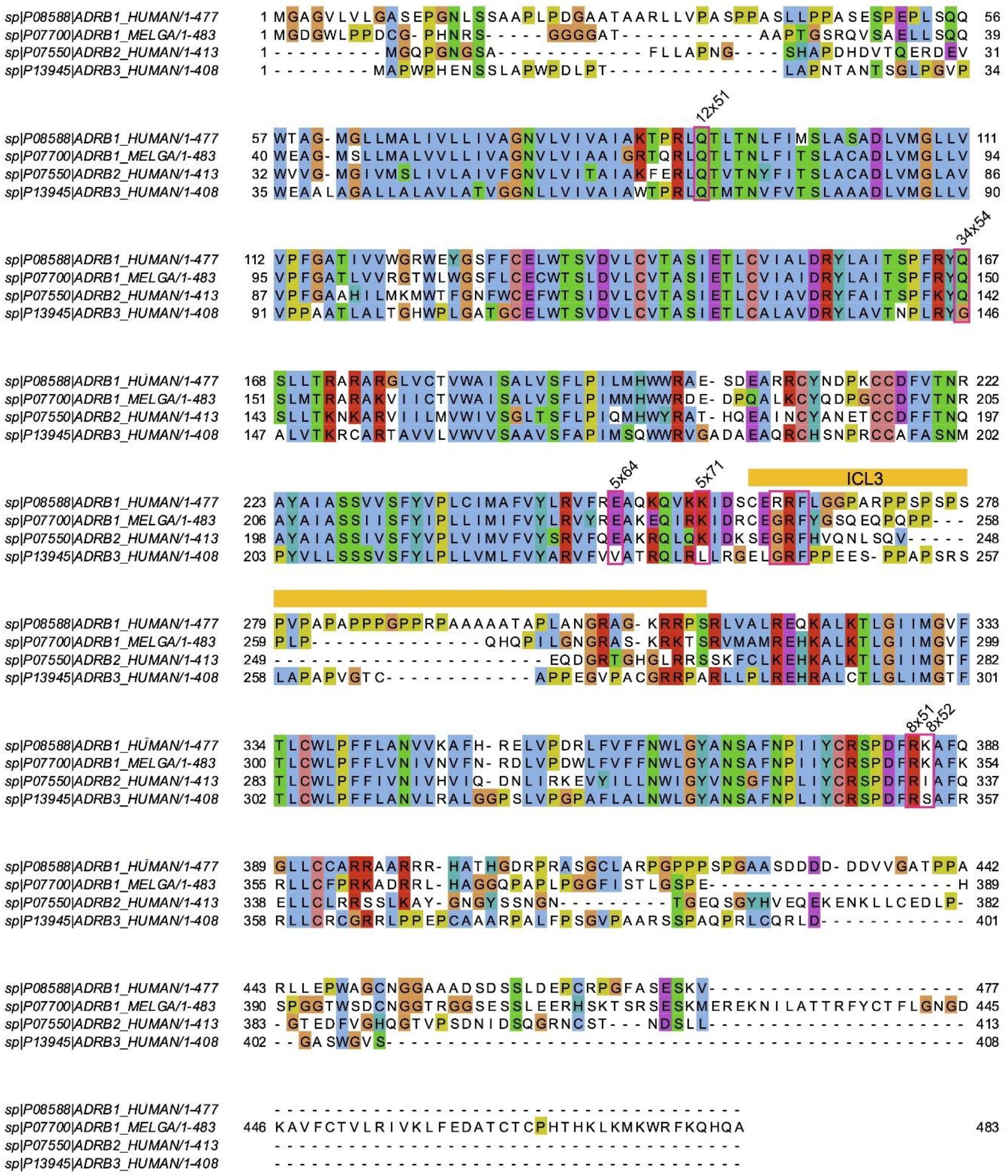

**Fig. S6. Sequence alignment of β-adrenergic receptors.** The sequence alignment was generated using UniProt database (The UniProt Consortium, Nucleic Acid Res 53 (2025)) and rendered using Jalview (Waterhouse AM et al., 2009). Residues contributing to the observed enhanced interface between the HFLβ1AR and $G_s$ are highlighted with pink rectangles and labeled with the revised Ballesteros-Weinstein system for class A GPCRs.[27, 28] The ICL3 region is highlighted in orange.

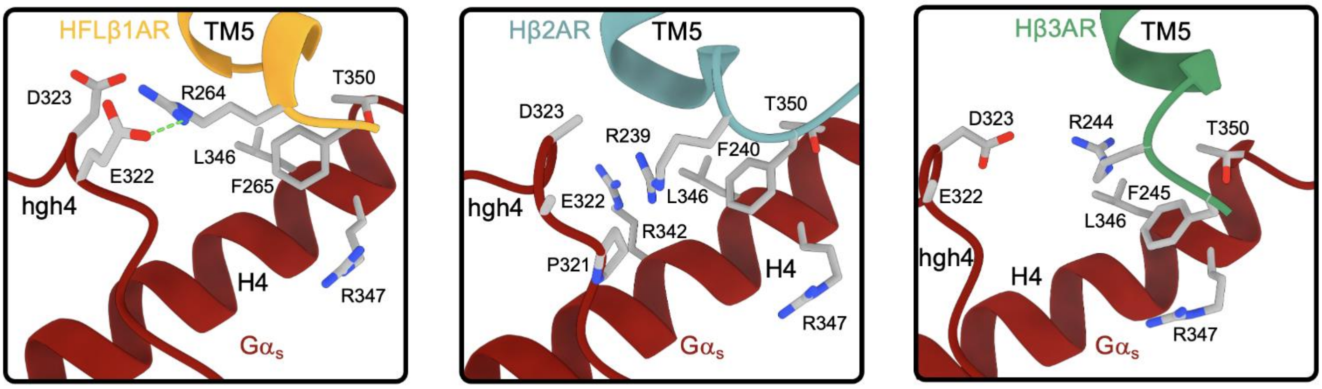

**Fig. S7. Comparison of ICL3 interactions with G$\alpha_s$ between the β-adrenergic receptors.** The receptors show similar hydrophobic interactions between $F^{ICL3}$ and H4 of G$\alpha_s$. $R^{ICL3}$ forms ionic interactions with hgh4 in the HFLβ1AR structure and is in close contact with residues in hgh4 or H4 in the β2AR-$G_s$ (PDBID 8GDZ) and β3AR-$G_s$ (PDBID 9IJE) structures.

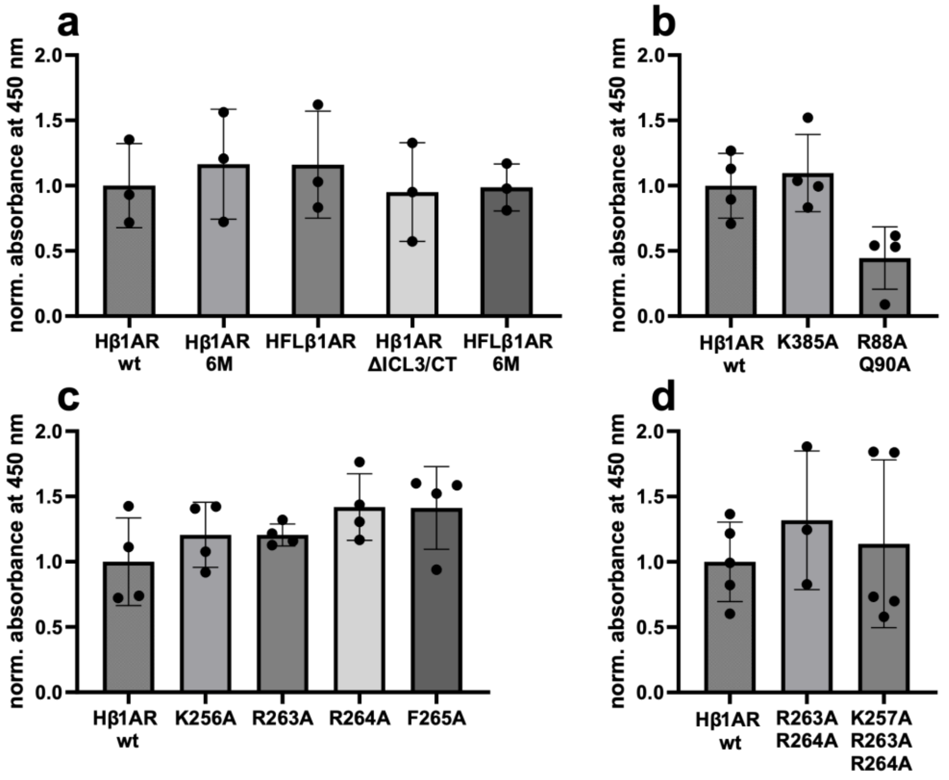

**Fig. S8. Surface expression levels of different Hβ1AR constructs and mutants.** Cell surface ELISA was performed on transiently transfected HEK293A cells. The empty vector control pcDNA was used for background subtraction and all values were normalized for each experiment to the average absorbance of Hβ1ARwt. n=3-5 mean ± SD. **a**. Expression levels of the various Hβ1AR constructs. **b-d**. Effect of mutations on the surface expression levels of Hβ1ARwt.

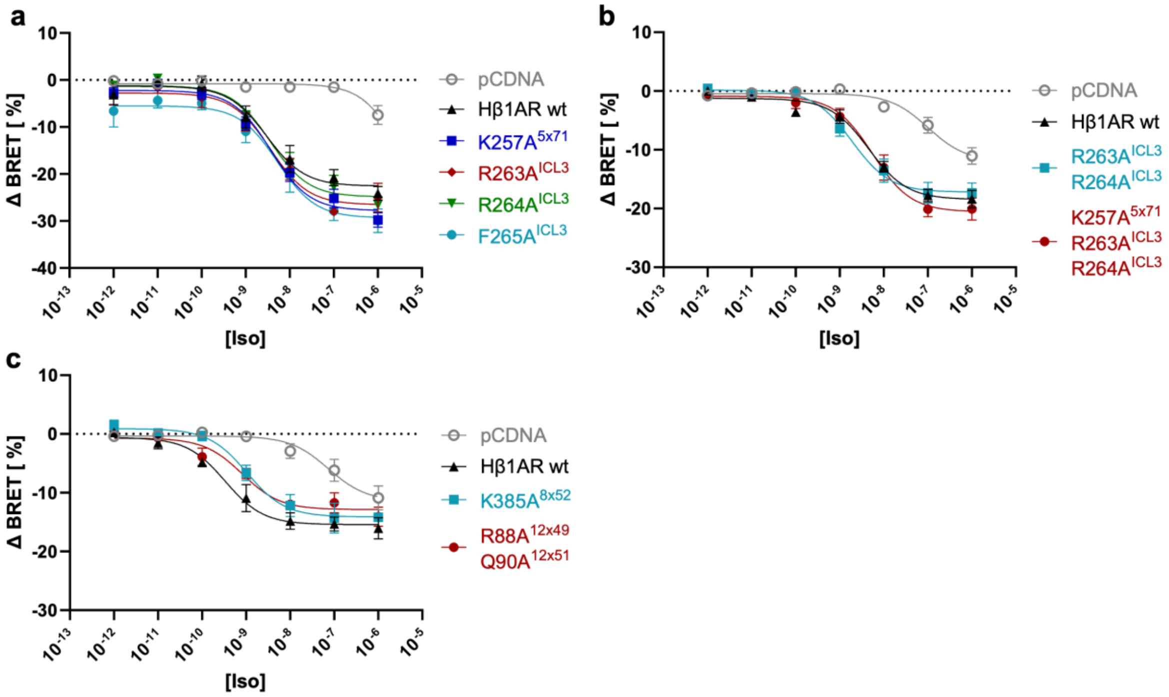

**Fig. S9. Influence of the identified and ICL3-mediated additional G protein-interacting residues on cAMP accumulation.** Signaling assays for point mutants in the extended interface between Hβ1ARwt and the heterotrimeric G protein. Signaling graphs represent the fit of grouped data ± SEM from three independent biological replicates. **a**. Single mutations of residues interacting with $G\alpha_s$. **b**. Combination of mutations of $G\alpha_s$-interacting residues. **c**. Combination of mutations of Gβ-interacting residues.

**Table S1. Cryo-EM data and model statistics**

| | (EMDB-19683) (PDB 8S2T) |
|---|---|
| **Data collection and processing** | |
| Magnification | 165,000x |
| Voltage (kV) | 300 |
| Electron exposure (e–/Å$^2$) | 65 |
| Exposure time (s) | 9 |
| Defocus range (µm) | -1.0 - -2.5 |
| Detector | K2 Summit |
| Pixel size (Å) | 0.8093 |
| Symmetry imposed | C1 |
| Initial particle images (no.) | 235,380 |
| Final particle images (no.) | 74,266 |
| Map resolution (Å) | 3.3 |
| FSC threshold | 0.143 |
| Map resolution range (Å) | 3.0 – 4.5 |
| **Refinement** | |
| Initial model used (PDB code) | 6E3Y and Colabfold |
| Map sharpening *B* factor (Å$^2$) | |
| Model composition | |
| Non-hydrogen atoms | 7,957 |
| Protein residues | 1,002 |
| Ligands | 1 |
| *Average B* factors (Å$^2$) | |
| Protein | 42.2 |
| Ligand | 37.6 |
| R.m.s. deviations | |
| Bond lengths (RMSZ) | 1.24 |
| Bond angles (RMSZ) | 1.02 |
| Validation | |
| MolProbity score | 0.87 |
| Clashscore | 1.2 |
| Poor rotamers (%) | 0.0 |
| Q-Score | 0.444 |
| Ramachandran plot | |
| Favored (%) | 99.3 |
| Allowed (%) | 0.7 |
| Disallowed (%) | 0.0 |
